# A *Vibrio* T6SS-mediated lethality in a marine animal model

**DOI:** 10.1101/2023.03.06.530577

**Authors:** Hadar Cohen, Motti Gerlic, Dor Salomon

## Abstract

Bacteria belonging to the genus *Vibrio* include many known and emerging pathogens. Horizontal gene transfer of pathogenicity islands is a major contributor to the emergence of new pathogenic *Vibrio* strains. In the current report, we use the brine shrimp *Artemia salina* as a model to show that the marine bacterium *Vibrio proteolyticus* uses a mobile type VI secretion system, T6SS3, to intoxicate eukaryotic hosts. Two T6SS3 effectors, which were previously shown to induce inflammasome-mediated pyroptotic cell death in mammalian phagocytic cells, contribute to this toxicity. Furthermore, we find a novel, widespread T6SS3 effector that also contributes to the lethality mediated by this system against *Artemia salina*. Therefore, our results reveal a T6SS that is shared among diverse vibrios and mediates host lethality, indicating that it can lead to the emergence of new pathogenic strains.

## Main Text

Global warming and the rise in ocean water temperatures are linked to the spread of Gram-negative bacteria of the genus *Vibrio* and their associated human diseases ^1^. Since vibrios are known for their ability to horizontally acquire new virulence traits ^2^, their continuing spread requires an in-depth investigation of their pan-genome virulence potential.

We previously reported that *Vibrio proteolyticus* (*Vpr*) strain NBRC 13287 (ATCC 15338) has three anti-eukaryotic determinants: a secreted pore-forming hemolysin (VPRH) that kills mammalian cells, and two type VI secretion systems (T6SS1 and T6SS3) that deliver anti-eukaryotic effector proteins into mammalian phagocytic cells ^3–5^. Here, we set out to investigate the virulence potential of these determinants in a marine animal model, *Artemia salina*.

Challenging *Artemia* nauplii with either wild-type or Δ*vprh Vpr* strains resulted in comparable high survival rates (**Fig 1A** and **Table S1**), suggesting that VPRH does not play a significant role during the infection of this marine animal. To investigate the virulence potential of the T6SSs, we used a strain in which we deleted the negative regulator *hns1*, leading to their hyper-activation ^4^. This Δ*vprh*/Δ*hns1* strain, which induced pyroptotic cell death in mammalian phagocytic cells ^4^, was significantly more lethal to the nauplii than the wild-type and Δ*vprh* strains (**Fig 1A** and **Table S1**); its hazard ratio was more than five times higher in comparison to the wild-type strain (**Fig. 1A**). Therefore, we posited that a *Vpr* T6SS is responsible for *Artemia* lethality.

**Figure 1.**
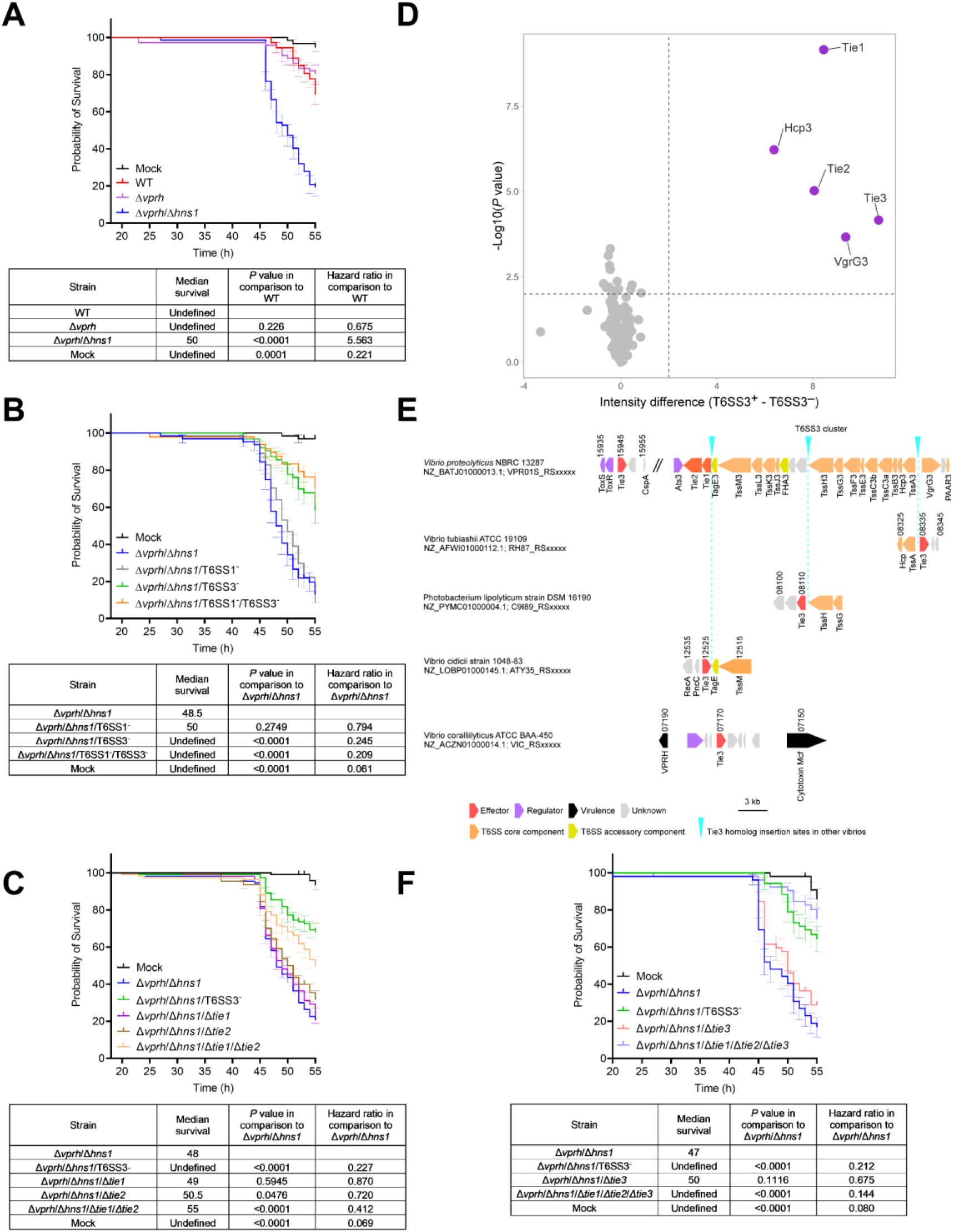
*Vpr* T6SS3 mediates lethality in *Artemia* nauplii. **(A-C, F).** *Artemia* nauplii were challenged with the indicated *Vpr* strains, and survival was assessed 20-55 hours post-infection (approximately 10^8^ bacteria were added to each well containing two nauplii). Comparisons were performed using the Log-rank (Mantel-Cox) test and Mantel-Haenszel hazard Ratio. Data are shown as the mean ± SE of at least three biological replicates. In (A), significant differences and hazard ratio are provided only for comparisons to the wild-type (WT) strain. In (B-C, F), significant differences and hazard ratio are provided only for comparisons to the Δ*vprh*/Δ*hns1* strain. A significant difference was considered as *P* < 0.05. **(D)** A volcano plot summarizing the comparative proteomics of proteins identified in the media of *Vpr* strains with an active (T6SS3^+^) or inactive (T6SS3^−^) T6SS3 and over-expressing Ats3 from a plasmid, using label-free quantification. The average difference in signal intensities between the T6SS3^+^ and T6SS3^−^ strains is plotted against the -Log10 of Student’s *t*-test *P* values (n = 3 biological replicates). Proteins that were significantly more abundant in the secretome of the T6SS3^+^ strain (difference in the average LFQ intensities > 2; *P* value < 0.03; with a minimum of 10 Razor unique peptides) are denoted in purple. **(E)** *Vpr* T6SS3 gene cluster and genomic neighborhoods of selected Tie3 homologs. Genes are represented by arrows indicating the direction of transcription. Locus tags are denoted above; encoded proteins and known domains are denoted below.

To determine which T6SS is responsible for the *Vpr*-mediated lethality, we challenged *Artemia* with strains in which we inactivated either T6SS1, T6SS3, or both (Δ*vprh*/Δ*hns1*/T6SS1^−^, Δ*vprh*/Δ*hns1*/T6SS3^−^, or Δ*vprh*/Δ*hns1*/T6SS1^−^/T6SS3^−^, respectively) by deleting essential components of the systems. Inactivation of T6SS1 had no significant effect on *Artemia* survival (**Fig. 1B** and **Table S2**). However, the inactivation of T6SS3 (either alone or in combination with T6SS1) resulted in significantly higher survival rates of *Artemia* and a four to five times lower hazard ratio compared to the Δ*vprh*/Δ*hns1* strain (**Fig. 1B** and **Table S2**). These results indicated that T6SS3, which is probably horizontally shared between pathogenic vibrios and induces pyroptotic cell death in mammalian phagocytic cells ^4^, mediates lethality in a marine animal.

We previously identified two *Vpr* T6SS3-secreted effectors, Tie1 and Tie2. We showed that these effectors are necessary and sufficient to induce pyroptotic cell death in mammalian phagocytic cells ^4^. Surprisingly, challenging *Artemia* nauplii with a strain in which we deleted both of these T6SS3 effectors (Δ*vprh*/Δ*hns1*/Δ*tie1*/Δ*tie2*) did not mirror the protection observed upon inactivation of T6SS3 as measured by survival, median survival time, and hazard ratio (**Fig. 1C** and **Table S3**). Although it was less lethal than its parental Δ*vprh*/Δ*hns1* strain, Δ*vprh*/Δ*hns1*/Δ*tie1*/Δ*tie2* was still more lethal than the Δ*vprh*/Δ*hns1*/T6SS3^−^ strain. This result led us to hypothesize that *Vpr* delivers additional effectors contributing to the T6SS3-mediated lethality in *Artemia*.

To test our hypothesis, we performed a comparative proteomics analysis to reveal the T6SS3 secretome using strains over-expressing the T6SS3-specific activator, Ats3 ^4^, from a plasmid. We identified five proteins that were preferentially secreted by the wild-type (T6SS3^+^) strain compared to the T6SS3^−^ strain: two secreted structural components of T6SS3 (Hcp3 and VgrG3), two known secreted effectors (Tie1 and Tie2), and a new hypothetical protein with no predicted activity or domain (WP_021706393.1) (**Fig. 1D** and **Dataset S1**). We predicted that the latter is also a T6SS3 effector; following the previous nomenclature of *Vpr* T6SS3 effectors ^4^, we named it Tie3. A secretion assay using custom-made antibodies confirmed that Tie3 is secreted in a T6SS3-dependent manner and is not required for T6SS3 activity (**Fig. S1**). Although Tie3 is encoded outside the main *Vpr* T6SS3 gene cluster, it is found next to the virulence regulators ToxRS ^6^. Notably, its homologs are often encoded within or flanking a T6SS3-like system, or next to known virulence factors (e.g., a VPRH homolog) (**Fig. 1E**).

The above results prompted us to investigate whether Tie3 plays a role in the T6SS3-mediated lethality in *Artemia*. Indeed, challenging *Artemia* with a strain in which all three effectors are deleted (Δ*vprh*/Δ*hns1*/Δ*tie1*/Δ*tie2*/Δ*tie3*) resulted in survival rates and a hazard ratio comparable to those observed for the Δ*vprh*/Δ*hns1*/T6SS3^−^ strain (**Fig. 1F** and **Table S4**). Notably, the deletion of *tie3* did not affect bacterial growth (**Fig. S2**). Therefore, Tie3 is a novel effector that contributes to the T6SS3-mediated lethality in *Artemia*, together with the previously described effectors Tie1 and Tie2. Future work will reveal its eukaryotic target and activity.

In conclusion, we found that although VPRH is lethal to mammalian cells, it does not appear to contribute to the virulence of *Vpr* in a marine animal model. Rather, a T6SS is responsible for *Vpr*-mediated lethality in *Artemia*, using effectors that also induce pyroptotic cell death in mammalian phagocytic cells. Since this virulent T6SS3 is horizontally shared between pathogenic vibrios and other marine bacteria ^4^, it may contribute to the emergence of new pathogenic strains.

## Acknowledgements

We thank members of the Gerlic and Salomon labs for helpful discussions, and the Smoler Proteomics Center at the Technion for performing and analyzing the mass spectrometry data. The research was supported by the Recanati Foundation (to MG and DS) and the Israel Science Foundation (grant number 2174/22 to MG).

