## Supplementary Information for "A *Vibrio* T6SS-mediated lethality in a marine animal model"

Materials and Methods

Supplementary Figures S1-S2

Supplementary Tables S1-S6

Supplementary References

### Materials and Methods

**Bacterial strains and media:** For a complete list of bacterial strains used in this study, see **Table S5**. *Vibrio proteolyticus* (*Vpr*) strains were grown in MLB (Lysogeny broth supplemented with NaCl to a final concentration of 3% [w/v]) or on MLB agar plates (supplemented with 1.5% [w/v] agar) at 30°C. Media were supplemented with kanamycin (250 µg/mL) or chloramphenicol (10 µg/mL) when appropriate to maintain plasmids. *Escherichia coli* (*E. coli*) strains were grown in 2xYT broth (1.6% [w/v] tryptone, 1% [w/v] yeast extract, and 0.5% [w/v] NaCl) or Lysogeny broth (LB) at 37°C. Media were supplemented with chloramphenicol (10 µg/mL) when appropriate to maintain plasmids. To induce the expression of genes from arabinose-inducible pBAD plasmids, 0.05% (w/v) L-arabinose was included in the media.

**Construction of plasmids and bacterial deletion strains:** For a complete list of plasmids used in this study, see **Table S6**. For in-frame deletions in *Vpr*, a $Cm^R$OriR6K suicide plasmid, pDM4 [1] containing 1 kb sequences upstream and downstream of each gene or region to be deleted were transformed into *E. coli* DH5α (λ-pir) by electroporation. The plasmids were then transferred into *Vpr* via conjugation. Trans-conjugants were selected on MLB agar plates containing chloramphenicol (10 µg/mL). The resulting trans-conjugants were grown on MLB agar plates containing 15% (w/v) sucrose for counter-selection and loss of the SacB-containing pDM4. Deletions were confirmed by PCR. For this work, the regions upstream and downstream of *tie3* were cloned into pDM4 using restriction digestion and ligation, as previously described [2].

***Artemia* infections:** *Artemia salina* eggs (Artemio Pur; JBL) were incubated in deionized distilled water supplemented with chloramphenicol (10 µg/mL), kanamycin (100 µg/mL), and ampicillin (100 µg/mL) for 1 hour at 28°C with constant rotation. The eggs were then washed three times with instant ocean (3.3% [w/v]; Aquarium Systems) and incubated overnight with constant rotation at 28°C. *Artemia* nauplii were transferred into sterile 48-well plates (two nauplii per well in 400 µL) and challenged with approximately $10^8$ colony forming units (CFU) of the indicated *Vpr* strains. The plates were then incubated at 28°C with 12 hours light and dark cycles. *Artemia* survival was scored at the indicated time points starting 20 hours post-infection. Each challenge was done in n≥8 wells. CFU of bacterial strains used for infection were determined at the time of infection (t = 0 hours) by counting colonies in tenfold serial dilutions spotted onto Marine Minimal Media (MMM) agar plates (1.5% [w/v] agar, 2% [w/v] NaCl, 0.4% [w/v] galactose, 5 mM $MgSO_4$, 7 mM $K_2SO_4$, 77 mM $K_2HPO_4$, 35 mM $KH_2PO_4$, and 2 mM $NH_4Cl$).

**Bacterial growth assays:** Overnight-grown cultures of *Vpr* were normalized to an $OD_{600}$ = 0.01 in MLB media and transferred to 96-well plates in triplicates (200 µL per well). The cultures were grown at 30°C in a BioTek EPOCH2 microplate reader with continuous shaking at 205 cpm. $OD_{600}$ readings were acquired every 10 minutes. Experiments were performed three times with similar results.

**Protein secretion assays:** *Vpr* strains were grown overnight in MLB media supplemented with antibiotics to maintain plasmids. The cultures were normalized to $OD_{600}$ = 0.18 in 5 mL MLB supplemented with appropriate antibiotics and 0.05% (w/v) L-arabinose to induce expression from the arabinose-inducible plasmids, and then grown for 5 hours at 30°C. For expression fractions (cells), 0.5 $OD_{600}$ units were collected, and cell pellets were resuspended in (2x) Tris-Glycine SDS sample buffer (Novex, Life Sciences). For secretion fractions (media), culture volumes equivalent to 10 $OD_{600}$ units were filtered (0.22 µm), and proteins were precipitated using deoxycholate and trichloroacetic acid [3]. Cold acetone was used to wash the protein precipitates twice. Then, protein precipitates were resuspended in 20 µL of 10 mM Tris-HCl pH = 8, followed by the addition of 20 µL of (2x) Tris-Glycine SDS sample buffer supplemented with 5% (v/v) β-mercaptoethanol. Next, 0.5 μL of 1 N NaOH were added to maintain a basic pH. The expression and secretion samples were boiled and then resolved on any-kD gradient Mini-PROTEAN Stain-Free precast gels (Bio-Rad). Expression and secretion of the indicated proteins were evaluated by immunoblotting with specific, custom-made antibodies against Hcp3 or Tie3 (polyclonal antibodies raised in rabbits against peptides: CQKHNYELEGGEIKD and CPVETPPHKDKTKRR, respectively; GenScript) used at 1:1000 dilution. Protein signals were visualized in a Fusion FX6 imaging system (Vilber Lourmat) using enhanced chemiluminescence (ECL) reagents. Equal loading was assessed using trihalo compounds' fluorescence of the immunoblot membrane.

**Mass spectrometry analyses:** Sample preparations for mass spectrometry were performed as described in the "Protein secretion assays" section. After the acetone wash step, samples were shipped to the Smoler Proteomics Center at the Technion, Israel, for analysis. Precipitated proteins were washed four times in 80% (v/v) cold acetone and incubated for 15 minutes at −20°C, followed by sonication. Ten μg of protein were reduced using DTT at 60ºC for 30 minutes. The proteins were then modified with 10 mM iodoacetamide in 100 mM ammonium bicarbonate for 30 minutes at room temperature in the dark. The proteins were digested overnight at 37ºC in 2 M urea and 25 mM ammonium bicarbonate with modified trypsin (Promega) at a 1:50 (M/M) enzyme-to-substrate ratio. An additional trypsinization step was performed for four hours in 1:100 enzyme-to-substrate ratio. The tryptic peptides were desalted using C18 tips (homemade stage tips), and then dried and re-suspended in 0.1% Formic acid. The resulting tryptic peptides were resolved by reverse-phase chromatography on 0.075 X 250-mm or 0.075 X 300-mm fused silica capillaries (J&W) packed with Reprosil reversed phase 358 material (Dr Maisch GmbH, Germany). The peptides were eluted with a linear 60-minute gradient of 5 to 28%, 15 minutes gradient of 28 to 95%, and 15 minutes at 95% acetonitrile with 0.1% formic acid in water at a flow rate of 0.15 μL/minute. Mass spectrometry was performed using a Q Exactive plus mass spectrometer (Thermo) in a positive mode using a repetitively full MS scan followed by high collision dissociation (HCD) of the 10 most dominant ions selected from the first MS scan. The mass spectrometry data were analyzed using MaxQuant software 1.5.2.8 for peak picking and identification using the Andromeda search engine [4] against the relevant *Vpr* strain from the Uniprot database with a mass tolerance of 6 ppm for the precursor masses and 20 ppm for the fragment ions. Oxidation on methionine was accepted as variable modifications, and carbamidomethyl on cysteine was accepted as static modifications. The minimal peptide length was set to seven amino acids; a maximum of two miscleavages was allowed. The data were quantified by label-free analysis using the same software. Peptide- and protein-level false discovery rates (FDRs) were filtered to 1% using target-decoy strategy. Statistical analysis of the identification and quantization results was done using Perseus 1.6.7.0 software [5]. Intensity data were transformed to log2. Values below the threshold were replaced with 21 (on the logarithmic scale), which corresponds to the lowest intensity that was detected. A Student's *t*-test with Permutation-based FDR (with 250 randomization, threshold value = 0.05) was performed. The mass spectrometry proteomics data have been deposited in the ProteomeXchange Consortium via the PRIDE [6].

**Identification of Tie3 homologues:** Tie3 (WP_021706393.1) homologs were identified using BLAST [7]. The genomic neighborhoods of selected homologs were manually examined, and representative regions with similar genetic composition were chosen for representation.

**Statistical analysis:** The data were analyzed using GraphPad prism 9. Unless otherwise indicated, the data are presented as the mean ± SE. Comparisons of survival curves were performed using Log-rank (Mantel-Cox) test and Mantel-Haenszel hazard Ratio. Statistical significance was considered at $P < 0.05$.

**Data Availability Statement:** The authors confirm that the data supporting the findings of this study are available within the article and its supplementary material. The mass spectrometry raw data files were deposited in ProteomeXchange.

### Supplementary Figures

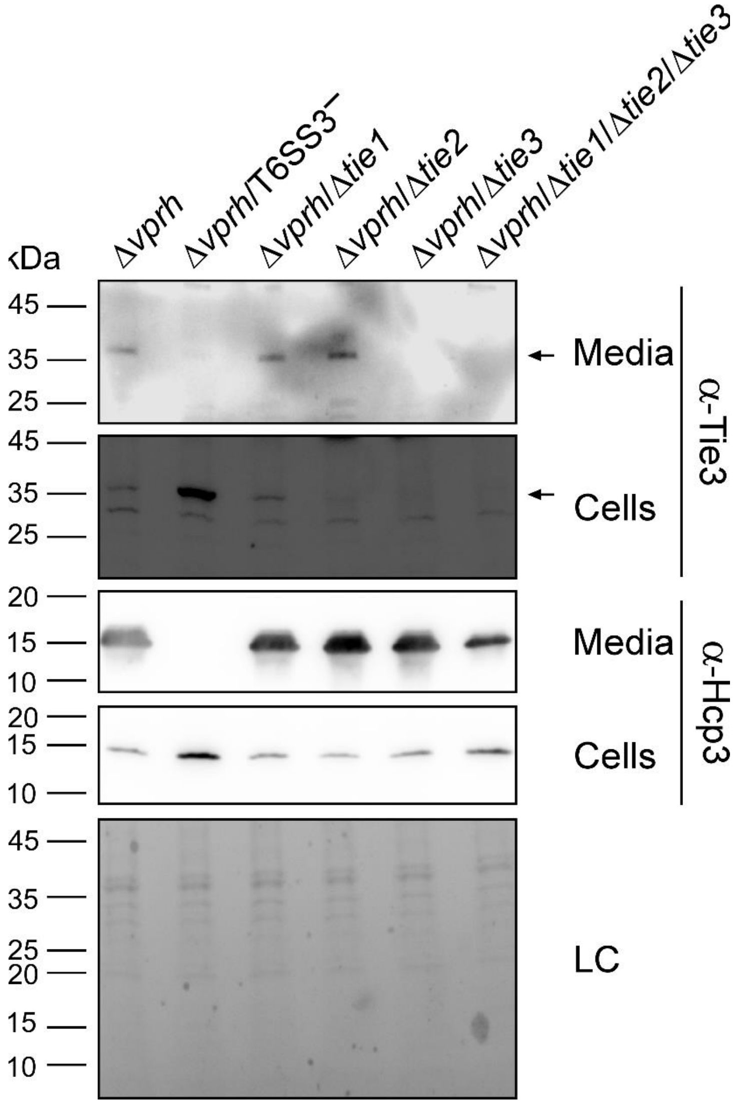


**Fig. S1. Tie3 is secreted in a T6SS3-dependent manner.** The expression (cells) and secretion (media) of Hcp3 and Tie3 from the indicated strains containing a plasmid for the arabinose-inducible expression of Ats3 were detected by immunoblotting using specific antibodies against Hcp3 and Tie3, respectively. Loading control (LC) is shown for total protein lysate. Arrows denote the expected size of Tie3. The data are representative of three independent experiment with similar results.

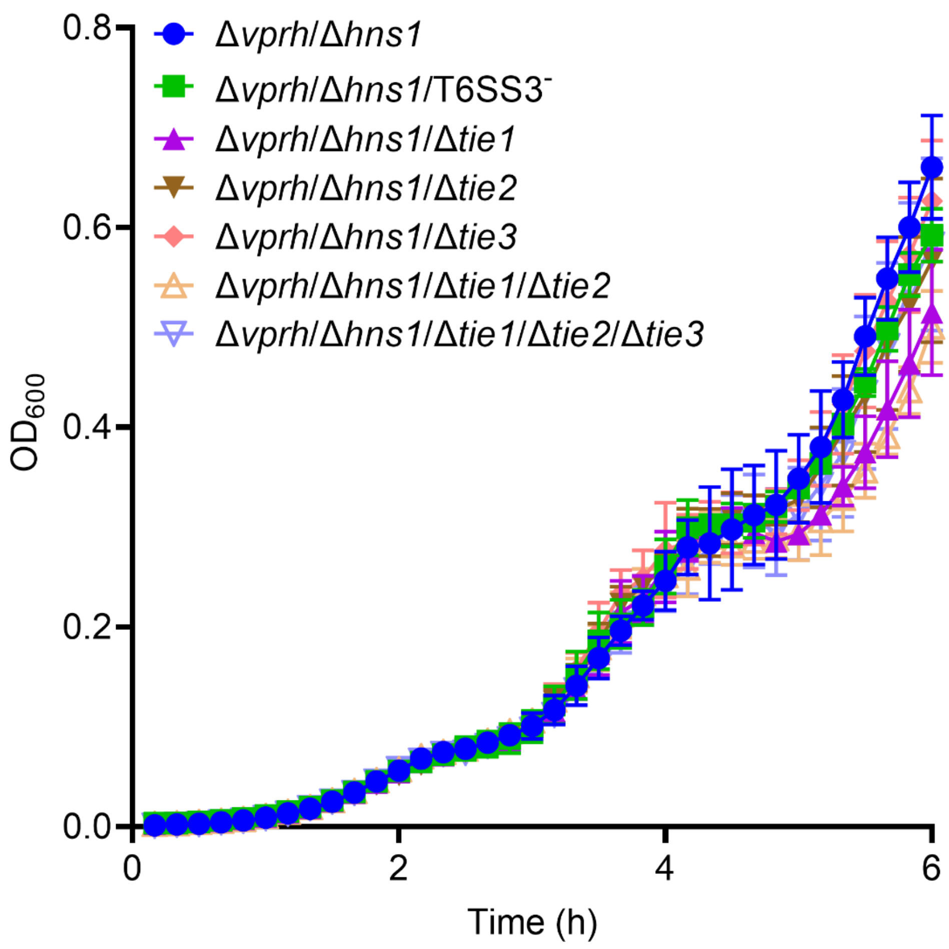


**Fig. S2. Deletion of *tie3* does not affect bacterial growth.** The growth of the indicated *Vpr* strains in MLB at 30°C measured as absorbance at 600 nm ($OD_{600}$). The data are a representative experiment out of three independent experiments.

### Supplementary Tables

**Table S1. Subjects at risk for Fig. 1A.**

| ***Vpr* strain** | **Time post-infection (h)** | | | | | | | |
|---|---|---|---|---|---|---|---|---|
| | **20** | **25** | **30** | **35** | **40** | **45** | **50** | **55** |
| **WT** | 72 | 72 | 72 | 72 | 72 | 72 | 70 | 56 |
| **Δ*vprh*** | 72 | 72 | 72 | 72 | 72 | 72 | 65 | 59 |
| **Δ*vprh*/Δ*hns1*** | 72 | 72 | 72 | 72 | 72 | 72 | 38 | 15 |
| **Mock** | 60 | 60 | 60 | 60 | 60 | 60 | 60 | 58 |

**Table S2. Subjects at risk for Fig. 1B.**

| *Vpr* strain | Time post-infection (h) | | | | | | | |
|---|---|---|---|---|---|---|---|---|
| | **20** | **25** | **30** | **35** | **40** | **45** | **50** | **55** |
| **Δ*vprh*/Δ*hns1*** | 64 | 64 | 64 | 63 | 63 | 60 | 25 | 9 |
| **Δ*vprh*/Δ*hns1*/T6SS1⁻** | 64 | 64 | 64 | 64 | 64 | 61 | 33 | 8 |
| **Δ*vprh*/Δ*hns1*/T6SS3⁻** | 63 | 63 | 63 | 63 | 63 | 61 | 53 | 22 |
| **Δ*vprh*/Δ*hns1*/T6SS1⁻/T6SS3⁻** | 48 | 48 | 48 | 48 | 48 | 47 | 42 | 13 |
| **Mock** | 64 | 64 | 64 | 64 | 64 | 64 | 63 | 31 |

**Table S3. Subjects at risk for Fig. 1C.**

| ***Vpr* strain** | **Time post-infection (h)** | | | | | | | |
|---|---|---|---|---|---|---|---|---|
| | **20** | **25** | **30** | **35** | **40** | **45** | **50** | **55** |
| **Δ*vprh*/Δ*hns1*** | 110 | 109 | 109 | 109 | 108 | 107 | 50 | 24 |
| **Δ*vprh*/Δ*hns1*/T6SS3⁻** | 110 | 110 | 110 | 110 | 110 | 109 | 90 | 60 |
| **Δ*vprh*/Δ*hns1*/Δ*tie1*** | 110 | 110 | 110 | 110 | 110 | 106 | 53 | 28 |
| **Δ*vprh*/Δ*hns1*/Δ*tie2*** | 110 | 110 | 110 | 110 | 105 | 103 | 59 | 30 |
| **Δ*vprh*/Δ*hns1*/Δ*tie1*/Δ*tie2*** | 110 | 108 | 108 | 108 | 107 | 106 | 78 | 47 |
| **Mock** | 110 | 110 | 110 | 110 | 110 | 110 | 110 | 85 |

**Table S4. Subjects at risk for Fig. 1F.**

| *Vpr* strain | Time post-infection (h) | | | | | | | |
|---|---|---|---|---|---|---|---|---|
| | **20** | **25** | **30** | **35** | **40** | **45** | **50** | **55** |
| **Δ*vprh*/Δ*hns1*** | 100 | 100 | 100 | 100 | 100 | 97 | 48 | 24 |
| **Δ*vprh*/Δ*hns1*/T6SS3$^-$** | 100 | 100 | 100 | 100 | 100 | 100 | 83 | 60 |
| **Δ*vprh*/Δ*hns1*/Δ*tie3*** | 52 | 52 | 52 | 52 | 52 | 50 | 30 | 11 |
| **Δ*vprh*/Δ*hns1*/Δ*tie1*/Δ*tie2*/Δ*tie3*** | 52 | 52 | 52 | 52 | 52 | 51 | 48 | 32 |
| **Mock** | 100 | 100 | 100 | 100 | 100 | 100 | 100 | 85 |

**Table S5. A list of bacterial strains used in this study.**

| Strain name | Genotype | Source |
|---|---|---|
| *Vibrio proteolyticus* NBRC 13287 (ATCC 15338) | Wild-type | ATCC |
| Δ*vprh* | *V. proteolyticus* ATCC 15338 Δ*vprh* | [8] |
| Δ*vprh*/T6SS3⁻ | *V. proteolyticus* ATCC 15338 Δ*vprh*/Δ*tssL3* | [9] |
| Δ*vprh*/Δ*hns1* | *V. proteolyticus* ATCC 15338 Δ*vprh*/Δ*hns1* | [9] |
| Δ*vprh*/Δ*hns1*/T6SS1⁻ | *V. proteolyticus* ATCC 15338 Δ*vprh*/Δ*hns1*/Δ*tssG1* | [9] |
| Δ*vprh*/Δ*hns1*/T6SS3⁻ | *V. proteolyticus* ATCC 15338 Δ*vprh*/Δ*hns1*/Δ*tssL3* | [9] |
| Δ*vprh*/Δ*hns1*/Δ*tie1* | *V. proteolyticus* ATCC 15338 Δ*vprh*/Δ*hns1*/Δ*tie1* | [9] |
| Δ*vprh*/Δ*hns1*/Δ*tie2* | *V. proteolyticus* ATCC 15338 Δ*vprh*/Δ*hns1*/Δ*tie2* | [9] |
| Δ*vprh*/Δ*hns1*/Δ*tie1*/Δ*tie2* | *V. proteolyticus* ATCC 15338 Δ*vprh*/Δ*hns1*/Δ*tie1*/Δ*tie2* | [9] |
| Δ*vprh*/Δ*hns1*/Δ*tie3* | *V. proteolyticus* ATCC 15338 Δ*vprh*/Δ*hns1*/Δ*tie3* | This study |
| Δ*vprh*/Δ*hns1*/Δ*tie1*/Δ*tie2*/Δ*tie3* | *V. proteolyticus* ATCC 15338 Δ*vprh*/Δ*hns1*/Δ*tie1*/Δ*tie2*/Δ*tie3* | This study |
| Δ*vprh*/Δ*tie1* | *V. proteolyticus* ATCC 15338 Δ*vprh*/Δ*tie1* | This study |
| Δ*vprh*/Δ*tie2* | *V. proteolyticus* ATCC 15338 Δ*vprh*/Δ*tie2* | This study |
| Δ*vprh*/ Δ*tie3* | *V. proteolyticus* ATCC 15338 Δ*vprh*/Δ*tie3* | This study |
| Δ*vprh*/Δ*tie1*/Δ*tie2*/Δ*tie3* | *V. proteolyticus* ATCC 15338 Δ*vprh*/Δ*tie1*/Δ*tie2*/Δ*tie3* | This study |

**Table S6. A list of plasmids used in this study.**

| Plasmid name | Description | Comments | Source |
|---|---|---|---|
| pDM4 | a $Cm^R$ and $ori_{R6K}$-containing suicide vector | Used as a backbone to construct plasmids for gene deletions in *Vibrio* | [1] |
| pDM4:*vprh* | pDM4 containing 1 kb upstream and 1 kb downstream of *vprh* in its MCS | Used to delete *vprh* in *V. proteolyticus* | [8] |
| pDM4:*hns1* | pDM4 containing 1 kb upstream and 1 kb downstream of *hns1* in its MCS | Used to delete *hns1* in *V. proteolyticus* | [9] |
| pDM4:*tssG1* | pDM4 containing 1 kb upstream and 1 kb downstream of *tssG1* in its MCS | Used to delete *tssG1* in *V. proteolyticus* | [10] |
| pDM4:*tssL3* | pDM4 containing 1 kb upstream and 1 kb downstream of *tssL3* in its MCS | Used to delete *tssL3* in *V. proteolyticus* | [9] |
| pDM4:*tie1* | pDM4 containing 1 kb upstream and 1 kb downstream of the region corresponding to nucleotides 485-584 of *tie1* in its MCS | Used to delete a 100 bp region within *tie1* in *V. proteolyticus* | [9] |
| pDM4:*tie2* | pDM4 containing 1 kb upstream and 1 kb downstream of *tie2* in its MCS | Used to delete *tie2* in *V. proteolyticus* | [9] |
| pDM4:*tie3* | pDM4 containing 1 kb upstream and 1 kb downstream of *tie3* in its MCS | Used to delete *tie3* in *V. proteolyticus* | This study |
| pAts3 | pBAD/Myc-$His^{Kan}$ containing the *ats3* ORF in its MCS, not fused to a C-terminal tag | Used for arabinose-inducible expression of Ats3 | [9] |

8 Ray A, Kinch LN, de Souza Santos M, Grishin N V., Orth K, Salomon D. Proteomics

Analysis Reveals Previously Uncharacterized Virulence Factors in Vibrio proteolyticus. *MBio* 2016; **7**: e01077-16.

9 Cohen H, Baram N, Fridman CM, Edry-Botzer L, Salomon D, Gerlic M. Post-phagocytosis activation of NLRP3 inflammasome by two novel T6SS effectors. *Elife* 2022; **11**: 1–25.

10 Ray A, Schwartz N, Souza Santos M, Zhang J, Orth K, Salomon D. Type VI secretion system MIX-effectors carry both antibacterial and anti-eukaryotic activities. *EMBO Rep* 2017; **18**: 1978–1990.
